# Autonomous drones are a viable tool for acoustic bat surveys

**DOI:** 10.1101/673772

**Authors:** Tom August, Tom Moore

## Abstract

Acoustic surveys of bats are currently limited by the detection range of ultrasound microphones. This makes it difficult to survey bats at height, over water, or in other hard to reach locations. Drones, also known as UAVs or UASs, are becoming more affordable and feature rich, resulting in more uptake in conservation, primarily for aerial imagery. In this paper we address current limitations to acoustic bat surveys by developing three autonomous drones for surveying bats; a plane, quadcopter and boat. All three are capable of moving autonomously from one waypoint to another while carrying a bat detector to record bats and are all low cost (under US$950) with large operational ranges that make them suited to aerial transect work.

Initial testing highlighted ultrasound noise generated by the drones as a major issue for recording bats from these systems. This was mitigated through iterative design of the microphone placement and the vehicles themselves. Subsequent testing in real world settings, in the presence of bats, demonstrated that bats could be recorded under autonomous navigation and that the ultrasound interference could be reduced to a negligible level.

Autonomous drones offer an exciting new tool in the tool box of bat workers and researchers. They allow us to study areas previously inaccessible on foot, as well as at heights that have previously been inaccessible. We also discuss current limitations to this technology, including legal considerations, hidden costs and potential impacts on bat behaviour.

## Introduction

Bats are key to ecosystem and human health as pollinators, pest controllers, and seed dispersers, amongst other roles (Kunz *et al* 2011). As a consequence, it is no surprise that a significant amount of time and money is spent studying bats’ distribution and behaviour in all corners of the globe. In the 1980s and 90s the emergence of commercially available bat detectors revolutionised the study of bats. Bat detectors are able to record the ultrasonic echolocaton calls that bats use to navigate in darkness. These recordings can be played back in audible sound in real-time, or analysed later on a computer, to determine species identity and individual behaviour. Bat detectors have since revolutionised our understanding of bats, yet there are still many limitations to their use. Principal among these is the detection range of bat detectors; approximately 10m to 20m (Limpers, 2004) though highly dependent on the species and detector used. This requires the surveyor to be within range of a bat to detect them and precludes the recording of bats at any significant height, above canopy, or in areas that are not easily accessible on foot such as over lakes, or in areas too dangerous to visit.

The height limitation of bat detectors has been addressed by some by placing bat detectors on masts and other tall structures (Collins and Jones 2009; Kalcounis *et al* 1999) or on helium-filled ballons (Fenton *et al* 1997; McCraken *et al* 2008). These studies have revealed greater bat activity above canopy than below (Kalcounis *et al* 1999), and even the presence of species at height (30m) that are not recorded at ground level (Collins and Jones 2009). This has profound implications not only for the study of bats’ ecology but also for activities, such as construction, that have impacts at height, notably wind turbines. While these previous studies have highlighted the need for a greater understanding of bat activity at height the methods used are limited. In the case of placement on tall structures, these surveys are static, only allowing recordings to be collected for a single point in the landscape, and require the presence of an existing structure or the temporary construction of one. Balloons are similarly constrained in their positioning, only able to sample in a vertical column and cannot be used in any significant amount of wind. There is therefore a need for a tool that can provide bat recordings at height with a similar freedom to that experienced when using a detector on the ground.

Some efforts have been made to record bats over water bodies such as lakes and rivers. In these studies detectors have either been mounted on floating platforms (Grindal *et al* 1999) or surveyors have used kayaks to take themselves and their recording equipment out onto the water (Lintott *et al* 2015). These methods are often the only way to get accurate recordings when the bankside vegetation prevents a clear view of the water or where the area of interest is greater than 10m-20m from the bankside.

We believe that recent developments in Unmanned Aerial Systems (UASs), also known as drones, have the potential to overcome some of these restrictions. Recent advances in UASs have made them cheaper and easier to use, with increased flight times, and as a result UASs have seen increased use in conservation research, for example to monitor wildlife populations using aerial imagery (Martin *et al* 2012; Hodgson *et al* 2016), and to map habitat at centimeter resolution (Habel *et al* 2016, Faye *et al* 2016). Almost all applications in conservation to date have focused on the collection of imagery of various types. To our knowledge there is only a single published study using a UAS for acoustic surveys of bats (Fu *et al* 2018). Fu *et al* present a UAS capable of flights of up to 18 minutes, costing $7729, which is only sufficient for static ‘point-of-interest’ surveys. In this paper we present UASs with much greater flight times which allows us to harness advances in software that enable UASs to fly autonomously, moving through a series of waypoints at set speed and altitude, either once or in a loop until instructed to land. This allows for precise, large scale and repeatable surveys with little human input. This software is not limited to UASs but can also be used to control unmanned ground vehicles or boats.

We present three, low-cost, unmanned systems - a plane, quadcopter and boat - designed specifically for the detection of bats using newly developed lightweight bat detectors. We highlight challenges in their design, construction and use, and show how these can be overcome. Finally, we will explore how these systems can be applied to further our understanding of the ecology of bats.

## Methods

We designed and built three types of unmanned autonomous drones to carry ultrasound recording equipment for detecting bats - a plane, quadcopter and boat. Planes and quadcopters currently commercially available are capable of flying pre-programmed routes, taking off and landing automatically and flying for over 30 minutes. However, in this fast-paced field this flight time is likely to improve over coming years. A quadcopter has four propellers and is able to hover like a helicopter, they are typically easier to fly than planes but for their cost have a shorter flight time and can lift less weight. Planes are harder to fly as they are constantly in motion to generate lift, however they are more energy efficient meaning that they have the ability to fly for longer and lift heavier loads. Boats benefit from expending no energy to stay stationary relative to the water, in contrast to both planes and quadcopters which constantly expend energy to stay airborne. Boats can also carry relatively large loads and since they only need to generate forward motion, not lift, they can run for much longer than UASs.

A lightweight bat detector was required to ensure agile control of the UASs and to maximise survey durations. Bat detectors are generally hand-held devices and are not built to be lightweight as a priority. It is important to have full spectrum recording so that all species present in the flight path of the UAS are recorded, and an audio trigger ensures that recordings were only made in the presence of bats, making the interpretation of data after the flight easier. We chose to use the Peersonic bat detector (www.peersonic.com) as it best met these needs, being 90g including battery, full spectrum and having a customisable trigger. We used the recorder without its outer casing to save on weight. In addition, we used a custom microphone that has reduced sensitivity to frequencies below 20 kHz.

Both UASs and the boat were tested for a) ability to move autonomously, b) ultrasound interference noise generated by electronics and motors, and c) ability to record bats in ‘real world’ settings. This process was iterative and numerous designs were built throughout the process, here we present the most advanced designs only. While there are many commercial UASs available we quickly moved to custom built UASs for a number of reasons. It allowed us to build UASs with custom components that reduced interference noise generated by the electronics, it allowed us to use open source software for autonomous navigation (Mission Planner, http://ardupilot.org/planner/), and it made the UASs more affordable. An overview of key features of each of the drones is given in Table 1, and a full parts list for each is available in the supplementary materials.

**Table 1.** Specification comparison between the three bat drones presented. Each drone as advantages and disadvantages and therefore the most appropriate drone is largely dependent on the application.

| **Feature** | **Plane** | **Quadcopter** | **Boat** |
| --- | --- | --- | --- |
| Weight | 3kg | 1.5kg | 2.5kg |
| Carrying capacity | 2kg | 1kg | 3kg |
| Maximum duration* | 90mins | 60mins | 120mins* |
| Build cost | £500 (\$674) | £700 (\$943) | £300 (\$404) |
| Time to build | 6 hrs | 15 hrs | 2 hrs |
| Maximum range* | 110km | 30km | 15km |
| Maximum speed | 30m/s | 20m/s | 15m/s |
| Minimum speed | 12 m/s | 0 m/s | 0 m/s |
| Waypoint accuracy | 5m | 5m | 5m |
| Easy of control | Very hard | Hard | Very easy |
| Launch requirements | 50m flat land | 2-5m clear space | Water access |

Prior to testing the UASs’ and boat’s ability to move autonomously, they were first tested under manual control to ensure they were safe. Autonomous control was tested by creating a transect of waypoints for the UAS to fly between in Mission Planner and then sending the vehicle out to navigate the waypoints. Success was assessed by comparing maps of waypoint routes and the actual route of the vehicle recorded in the onboard GPS logs.

Ultrasound interference produced by the UASs and boat was measured using the bat detector and BatSound software (www.batsound.com). Recordings were made at multiple locations on and around each vehicle to quantify the interference produce. These were used to optmise the components used in the final designs and to locate the best position for housing the detector.

Performance in ‘real world’ scenarios was tested by sending drones on autonomous transects at night in the presence of bats. The quadcopter and boat were operated at approximately 2 m/s, similar to walking speed, while the plane was operated at 12 m/s. UASs were kept at a minimum of 10 meters above the tallest vegetation, and so out of the flight path of most bats. Recordings were analysed and the relative amplitude of bat calls and interference from the vehicle was assessed.

## Results

Three vehicles were built, each capable of autonomous movement and each costing less than £700 ($943), excluding the cost of the bat detector (£274 / $353). Maximum flight times for the quadcopter (. 1a & 1b) and plane (fig. 1c) were estimated at 60 and 90 mins respectively, allowing for a total flight distance of approximately 30 km and 110 km. The boat (fig. 1d) was found to be much more energy efficient, and could achieve an estimated maximum duration of 120 mins at an average of 2 m/s, thereby covering a distance of circa 15 km (table 1).

**Figure 1.**
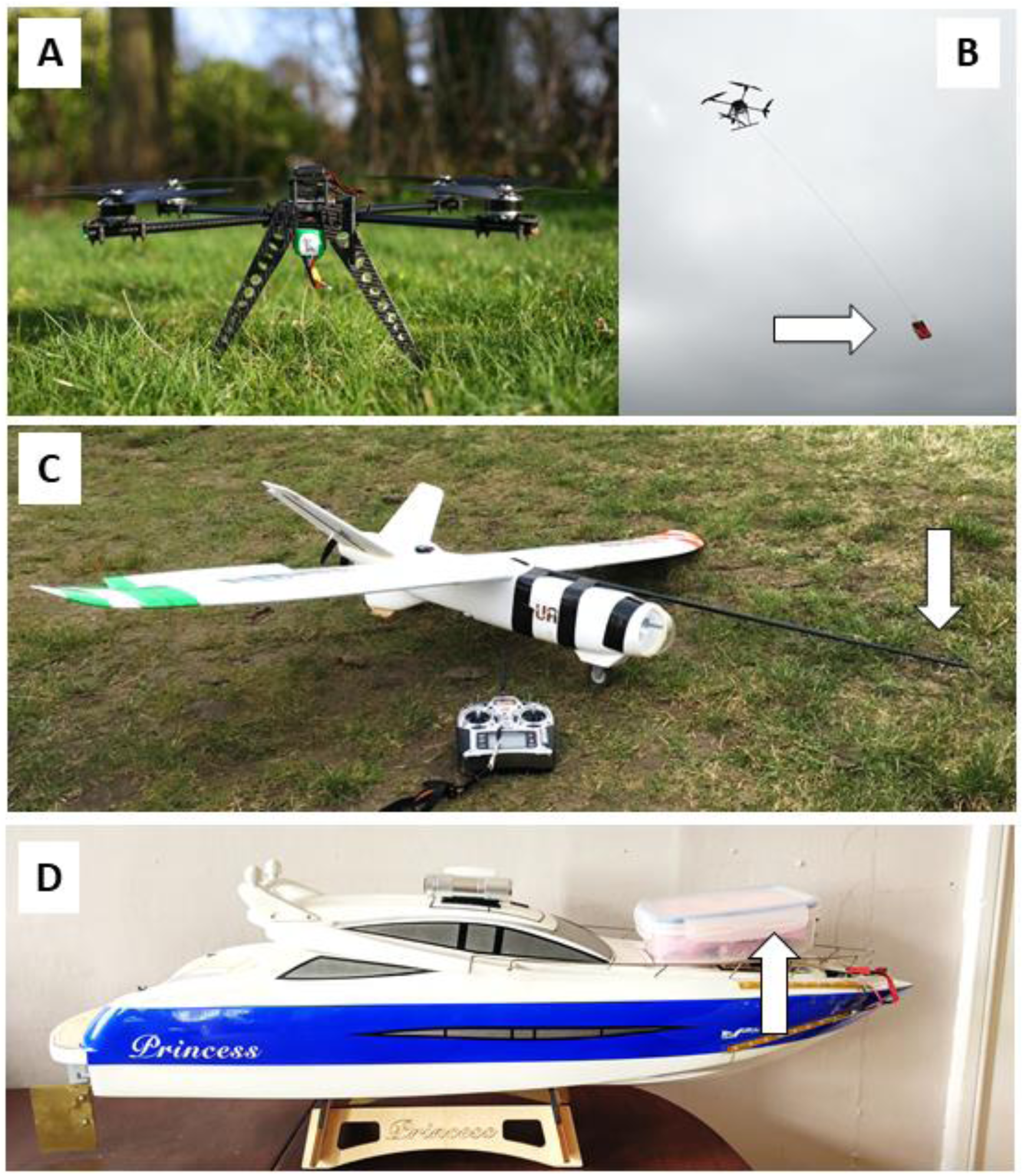
Images show the three drones designed and used to record bats ultrasound calls. A) Shows the quadcopter in profile B) shows the quadcopter in flight, the detector is suspended on 3m of nylon cord C) shows the plane, the microphone is attached to a rod on the front of the plane (arrow), while the body of the detector is house internally. D) shows the boat, the detector is housed in a waterproof box on the front of the boat (arrow), a hole in the front of the box, protected by water resistant gause, allows sound to enter the microphone.

### Autonomous movement

Both plane and quadcopter were able to safely and accurately navigate the waypoints, however given the quadcopter’s ability to hover it was able to follow the waypoints with greater accuracy than the plane. The boat was able to navigate waypoints along a river with high accuracy however the large turning circle of the boat model used made turning around difficult. The current boat would work in a larger water body such as a lake however at present is not suitable for use in rivers. We suggest that changing the model of boat to one with the rudder directly behind the propeller would easily overcome this issue by allowing the boat to achieve a much tighter turning circle.

### Ultrasound interference

Tests of the ultrasound produced by a number of UASs prior to the two aerial systems presented here revealed that ultrasound produced by the motors and propellers would significantly reduce the ability of a bat detector to record bats, and may affect the behaviour of bats in proximity. This challenge was addressed from two directions: firstly the UASs were designed to reduce the amount of noise produced by the motors and propellers and secondly, the distance between the propeller and the microphone was increased.

In fig. 2 we compare the ultrasound produced by our first two UAS designs (quadcopter – phantom 3, and plane – Bix3) with the final two designs presented in this paper. Through design (choice of propellers and motors, and positioning of the microphone) we were able to reduce the maximum decibels of noise from −18db to −44db and −20db to −65db for the plane and quadcopter respectively. The peak noise for both systems was at 25-30 KHz. This is a significant improvement, bearing in mind that decibels are on a logarithmic scale. Improvements were in part made by separating the microphone from the source of the ultrasound. This takes advantage of the physical properties of ultrasound, which attenuates far more quickly over distance than lower frequency audible sound. Separating the microphone by 3m on the quad was sufficient to reduce the noise to a minimal level that was not likely to interfere with the recording of bats. This separation was achieved by suspending the detector under the UASs using nylon cord. This distance did not cause any significant difficulty in flight, take-off, or landing. The plane design was chosen because the propeller is at the back of the aircraft; this allows a microphone at the front of the aircraft to be at a greater distance from the source of the ultrasound and in ‘clean’ air, undisturbed by the propeller. This advantage was further improved by placing the microphone on a rod extending from the front of the plane, attached to the top of the fuselage (fig 1d). This rod did not impact on the airworthiness of the aircraft.

**Figure 2.**
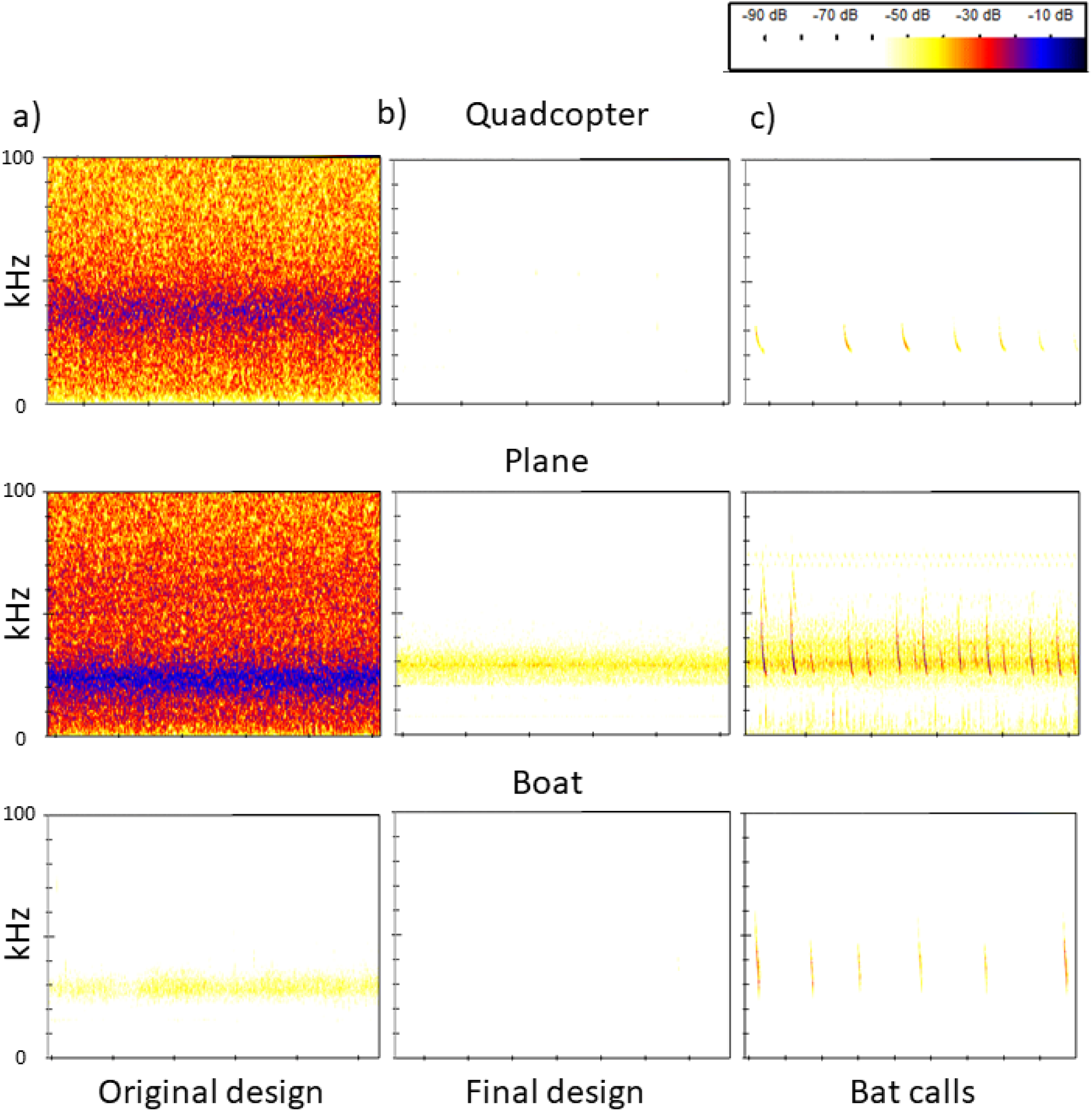
The noise produced by each drone (top row – quadcopter, middle row - plane, bottom row - boat) was greatly reduced when comparing the original designs (first column) to the final designs (middle column). For both UASs the original design had the detector attached to the body of the aircraft, close to the propeller. Testing in ‘real-world’ scenarios show that bats can be recorded from all three drones (right column).

### Real-world testing

All three systems were tested in ‘real world’ scenarios, at sites where bats were known to be active from ground-based detector surveys. Bats were recorded on all systems allowing an assessment of signal to noise ratios.

The boat produces the least interference noise of all three systems as, at slow speeds the propeller is very quiet. As a result, the signal to noise ratio is very high and bats call recordings are equivalent to those that would be recorded from a hand-held detector (fig. 2, c) boat). The quadcopter also had a good signal to noise ratio; the bat calls recorded can be easily seen (fig. 2, c) quadcopter), even those of *Myotis* species which cross the frequency of noise from the quadcopter (30-40 kHz). The signal to noise ratio was worst for the plane, however even with this system bat calls can be clearly seen above the noise generated by the motor and propeller.

## Discussion

We have described three unmanned systems capable of recording bats via onboard, lightweight, bat detectors. These systems overcome the challenge presented by ultrasonic noise generated by the propellers and take advantage of new battery and motor technology to produce flight times that allow for meaningful surveys. The cost of the drones presented is low enough to be accessible to conservation organisations or interested individuals. The cost is kept low by building the drones from components, using a low-cost bat detector, and using software that is free to use. The flight times of the UASs and the operational time of the boat allow for large scale surveys, going beyond what has previously been achieved (Fu *et al* 2018).

While foot surveys for bats, using hand held bat detectors, will continue to be the most effective means of collecting data in the majority of situations, bat detecting UASs and boats offer new opportunities, allowing us to study bats in previously inaccessible places and opening up new avenues of research. We see a number of opportunities for the application of drones, notably; at height, over water, at a large scale, and in inaccessible areas.

Bats are known to fly at height, such as over tree canopies, at heights greater than 30m (Collins and Jones 2009; Kalcounis *et al* 1999) and various studies have detected bat feeding activity at heights from 400-900m (Fenton *et al* 1997, McCracken *et al* 2008). The limitations of handheld detectors mean that currently there are few practical ways to survey bats at this height. As a consequence, we are currently at risk of making poor conservation choices when planning development at height. In particular this would apply to wind turbines, which are known to have negative impacts on bats (Baerwald *et al* 2008), and other tall structures. The use of UASs at height could also be used to better understand the foraging behaviour of bats at height (Fenton *et al* 1997, McCracken *et al* 2008) as well as investigate evidence of migration at height.

Bat surveys over water to date have revealed impacts of pollution and eutrophication (Salvarina *et al* 2016). The application of autonomous boats or UASs could further our understanding of the foraging behaviour of bats over water and explore the predictor variables of foraging activity such as pollution levels, bank side vegetation and human activity. These vehicles could collect data from across water bodies with a high level of repeatability, both during a single visit and across multiple visits to a site.

The flight duration and distance, most notably for UAS planes, offers opportunities for large scale, repeatable surveys of bats that would simply not be possible until present. Performing large scale surveys with UASs comes with additional considerations, most notably legal considerations. The legal restrictions for UAS flights is variable by country and is under constant review, however long-distance flights beyond the line of sight of the controller are in general heavily restricted. This means that the potential for long distance surveys may only be realised in countries with accommodating legislation.

Areas that are inaccessible on foot are currently very difficult to survey for bats. Examples include dense woodland, wetland and swamp habitats, and steep terrain, including cliffs. UASs offer a tool for rapidly assessing these habitats for the presence of bats and an opportunity to learn about the ecology of bats in these locations.

While we have highlighted a long list of opportunities offered by autonomous vehicles to bat researchers there are a number of limitations that must be considered when using them for monitoring bats. In particular it is important to consider the impacts on bat behaviour, flights in areas of variable terrain height and unforeseen costs.

The vehicles presented all create some ultrasound, though this has been designed to be minimal, and carry lights so that they can be seen and controlled by a pilot if manual control is required. We must understand the effects that this has on bats before we can reliably interpret the results of UAS ultrasound surveys. Fu *et al* noted, “*minimal impact on bat behaviour*”, when using their multi-rotor UAS which is considerably larger and louder than the UASs presented here. In our experience, bats did not appear disturbed by the UASs we have presented in this paper, however a study dedicated to testing this is required.

Our UASs are not aware of objects around them, meaning they are prone to collide with trees if they are not programmed to fly at a height guaranteed to be above tree canopy. As a consequence, flights are likely to be conservatively high where tree height or terrain height is heterogeneous, potentially too far from foraging bats to get useful recordings. New technologies are available that allow UASs to ‘see’ objects around them either using infrared, ultrasound or lasers. These could be used to allow UASs to control their altitude so that their relative height above the canopy or ground remains constant throughout a survey.

Autonomous vehicles represent a potential cost and time saving when compared to bat surveys made on land by foot, or on water by canoe. However, it is important to include training, maintenance and set-up times in efficiency estimates. In some countries, such as the UK, a course and operations manual are required to undertake commercial work with a UAS, this is costly both financially and in terms of time. Hidden costs such as maintenance, replacement parts and insurance should also be considered.

In conclusion, we have presented three autonomous drones that can be used for surveying bats. These offer bat researchers and surveyors tools to survey bats using large-scale, highly repeatable transects at height or on water. While these tools offer new opportunities and potential cost savings, current limitations include legal restrictions, training, and collision risks in heterogeneous landscapes.

## Supporting information

Supplementary Material - Parts lists

## Acknowledgements

We are grateful for the support of a range of individuals and organisations who have given time, resources and advice in support of this work. Thanks to Nigel Fisher, Ross Wingfield, Peersonic, 4D, Heliguy, easycomposites, Gliders Distribution, packaging2buy and members of the UK bat workers Facebook group in particular.

