## Supplementary Material - Parts lists for "Autonomous drones are a viable tool for acoustic bat surveys"

Quadcopter:

| **Type** | **Model** | **Source** | **Number required** |
| --- | --- | --- | --- |
| Motor | MN3510 360KV | - | 4 |
| Propeller | 15x5 CF | - | 4 |
| ESC | X-rotor 15A OPTO | - | 4 |
| Flight controller | 3DR Pixhawk 1 | <https://3dr.com/> | 3DR Pixhawk 1 |
| Battery | 4Ah 4S 10C Lipo | - | 1 |
| Frame | 12mm OD 10mm ID woven carbon tubes & 1.5mm 3K carbon plate | - | 1 |
| LED lights | 5mm red + green 12V | - | 4 |
| GPS | 3DR uBlox LEA-6H | <https://3dr.com/> | 1 |
| Telemetry radio | 3DR 433MHz telemetry | <https://3dr.com/> | 1 |
| Receiver | Spektrum satellite receiver | [www.ebay.co.uk](http://www.ebay.co.uk/) | 1 |
| Pixhawk 6S Power Module *2 | Generic 6-10S | [www.ebay.co.uk](http://www.ebay.co.uk/) | 1 |

Plane:

| **Part type** | **Part** | **Source** | **Number required** |
| --- | --- | --- | --- |
| Airframe | X-UAV Talon | [banggood.com](http://www.banggood.com/) | 1 |
| Airframe material | EPO | - | 1 |
| Flight controller | 3DR Pixhawk 1 | <https://3dr.com/> | 1 |
| GPS | 3DR uBlox LEA-6H | <https://3dr.com/> | 1 |
| Telemetry radio | 3DR 433MHz telemetry | <https://3dr.com/> | 1 |
| Receiver | Spektrum satellite receiver | [www.ebay.co.uk](http://www.ebay.co.uk/) | 1 |
| Motor | T-motor 4014 400KV | [www.heliguy.com](https://www.heliguy.com/) | 1 |
| Propeller | Aeronaut 15x6 CAM Carbon folding | [www.gliders.uk.com](http://www.gliders.uk.com/) | 1 |
| ESC | Overlander XP2 60A brushless w/5A SBEC | [www.gliders.uk.com](http://www.gliders.uk.com/) | 1 |
| Battery | Multistar 8000mAh 6S LiPo | [hobbyking.com](https://hobbyking.com/en_us) | 1 |
| Wing servos | Hitec HS-82MG | [www.servoshop.co.uk](http://www.servoshop.co.uk/) | 4 |
| Tail servos | Turnigy TGY-50090M Analog | [hobbyking.com](https://hobbyking.com/en_us) | 1 |
| Pixhawk 6S Power Module *2 | Generic 6-10S | [www.ebay.co.uk](http://www.ebay.co.uk/) | 1 |

Boat:

| **Part type** | **Part** | **Source** | **Number required** |
| --- | --- | --- | --- |
| Hull | Princess Hobby Boat | [www.hobbyking.com](http://www.hobbyking.com) | 1 |
| ESC | Turnigy marine 180A BEC watercooled ESC | [www.hobbyking.com](http://www.hobbyking.com) | 1 |
| Flight controller | 3DR Pixhawk 1 | <https://3dr.com/> | 1 |
| GPS | 3DR uBlox LEA-6H | <https://3dr.com/> | 1 |
| Telemetry radio | 3DR 433MHz telemetry | <https://3dr.com/> | 1 |
| Receiver | Spektrum satellite receiver | [www.ebay.co.uk](http://www.ebay.co.uk/) | 1 |
| Rudder servo | Spektrum S6180 digital servo | [www.spektrumrc.com](http://www.spektrumrc.com) | 1 |
| Rudder increased surface area | 2mm brass plate | Local hobby shop | 1 |
| Battery | Turnigy 6400mAh 3S Lipo | [www.hobbyking.com](http://www.hobbyking.com) | 1 |
| Pixhawk 6S Power Module *2 | Generic 6-10S | [www.ebay.co.uk](http://www.ebay.co.uk/) | 1 |
